# Proteogenomic annotation of the Chinese hamster reveals extensive novel translation events and endogenous retroviral elements

**DOI:** 10.1101/468181

**Authors:** Shangzhong Li, Seong Won Cha, Kelly Hefner, Deniz Baycin Hizal, Michael Bowen, Raghothama Chaerkady, Robert N. Cole, Vijay Tejwani, Prashant Kaushik, Michael Henry, Paula Meleady, Susan T. Sharfstein, Michael J. Betenbaugh, Vineet Bafna, Nathan E. Lewis

**Affiliations:** Department of Bioengineering, University of California, San Diego, La Jolla, CA, USA; Novo Nordisk Foundation Center for Biosustainability, University of California, San Diego, La Jolla, CA, USA; Department of Electrical and Computer Engineering, University of California, San Diego, La Jolla, CA, USA; Chemical and Biomolecular Engineering, Johns Hopkins University, Baltimore, Maryland, USA; Antibody Discovery and Protein Engineering, MedImmune LLC, Gaithersburg, Maryland USA; The Mass Spectrometry Core, Johns Hopkins School of Medicine, Baltimore, MD, 21205; Colleges of Nanoscale Science and Engineering, SUNY Polytechnic Institute, Albany, NY, USA; National Institute for Cellular Biotechnology, Dublin City University, Dublin 9, Ireland; Department of Computer Science and Engineering, University of California, San Diego, La Jolla, CA, USA; Department of Pediatrics, University of California, San Diego, La Jolla, CA USA

**Keywords:** Chinese hamster, genome annotation, proteogenomics, endogenous retrovirus

## Abstract

A high quality genome annotation greatly facilitates successful cell line engineering. Standard draft genome annotation pipelines are based largely on *de novo* gene prediction, homology, and RNA-Seq data. However, draft annotations can suffer from incorrectly predictions of translated sequence, incorrect splice isoforms and missing genes. Here we generated a draft annotation for the newly assembled Chinese hamster genome and used RNA-Seq, proteomics, and Ribo-Seq to experimentally annotate the genome. We identified 4,333 new proteins compared to the hamster RefSeq protein annotation and 2,503 novel translational events (e.g., alternative splices, mutations, novel splices). Finally, we used this pipeline to identify the source of translated retroviruses contaminating recombinant products from Chinese hamster ovary (CHO) cell lines, including 131 type-C retroviruses, thus enabling future efforts to eliminate retroviruses by reducing the costs incurred with retroviral particle clearance. In summary, the improved annotation provides a more accurate platform for guiding CHO cell line engineering, including facilitating the interpretation of omics data, defining of cellular pathways, and engineering of complex phenotypes.

## Introduction

Chinese hamster ovary (CHO) cells are the primary workhorse for therapeutic protein production(Golabgir et al., 2016). Sequencing and assembly of the CHO and Chinese hamster genomes(Brinkrolf et al., 2013a; Lewis et al., 2013a; Xu et al., 2011) have enabled improvement in protein production using genetic engineering and in cell line process optimization using omics technologies(Kuo et al., 2017; Stolfa et al., 2018). A recent effort greatly improved the reference Chinese hamster genome assembly by combining Pacific Biosciences Single Molecule Real Time (SMRT) and short-read Illumina sequencing data, thus reducing the number of scaffolds by 28-fold and filling 95% of the sequence gaps(Rupp et al., 2018). Despite these great improvements in the assembly, the current genome annotation was based primarily on *ab initio* prediction, protein homology, ESTs, and limited publicly available transcriptomic data. However, these pipelines have difficulties in translation confirmation, splice form detection, and complete novel gene identification(N. Castellana & Bafna, 2010). To improve cell line engineering success, an accurate genome annotation is necessary.

Proteogenomics provides a way to address such challenges by integrating mass spectrometry-based proteomics, RNA-Seq and genomic data. For example, peptides can be identified by mapping tandem mass spectra to protein databases derived from RNA-Seq and genome annotation. The peptides are then used to update the annotation with novel coding regions and splice sites. Proteogenomics was first applied to *Mycoplasma pneumoniae(Jaffe, Berg, & Church, 2004)* to identify new and extended open reading frames (ORFs) and remove low quality gene models. It has also been applied to many eukaryotes including plants(N. E. Castellana et al., 2008), yeast(Yagoub et al., 2015), and human(M.-S. Kim et al., 2014). In addition to improving annotations, the proteomic data can also identify mutations (e.g., in cancer(Woo et al., 2015) and post translational modifications(Cesnik, Shortreed, Sheynkman, Frey, & Smith, 2016; Kaushik, Henry, Clynes, & Meleady, 2018)).

In addition to proteomics data, Ribo-Seq (data from sequencing ribosome-protected coding reactions at single nucleotide resolution(Ingolia, Ghaemmaghami, Newman, & Weissman, 2009)), provides a global view of actively translated mRNAs, and thus has been utilized to predict ORFs and translation frames for proteins(Calviello & Ohler, 2017) and to identify additional predicted proteins for proteogenomics(Crappé et al., 2015). Together transcriptomic, Ribo-Seq, and proteomic data can be invaluable for refining annotation about proteins.

To obtain a data-supported refinement of the Chinese hamster genome annotation, here we integrated proteomics, RNA-Seq, and Ribo-Seq to verify coding regions, update gene models, identify novel translated genes, and verify protein-coding variants in different CHO cell lines **(Figure 1)**. To further demonstrate the increased value of this resource, we investigated the challenge associated with the Food and Drug Administration (FDA) requirement to ensure that viral particles (particularly endogenous retroviral particles) are eliminated from the therapeutic protein product, which contributes to the high costs in bioprocessing(Strauss et al., 2009). Specifically, we identified all translated retrovirus particles in CHO cells, including previously unannotated translated loci, thus providing potential knockout targets to increase drug purity and reduce demands on viral clearance. This proteogenomic resource will be invaluable for future efforts to study and engineer CHO cells for bioprocessing.

**Figure 1:**
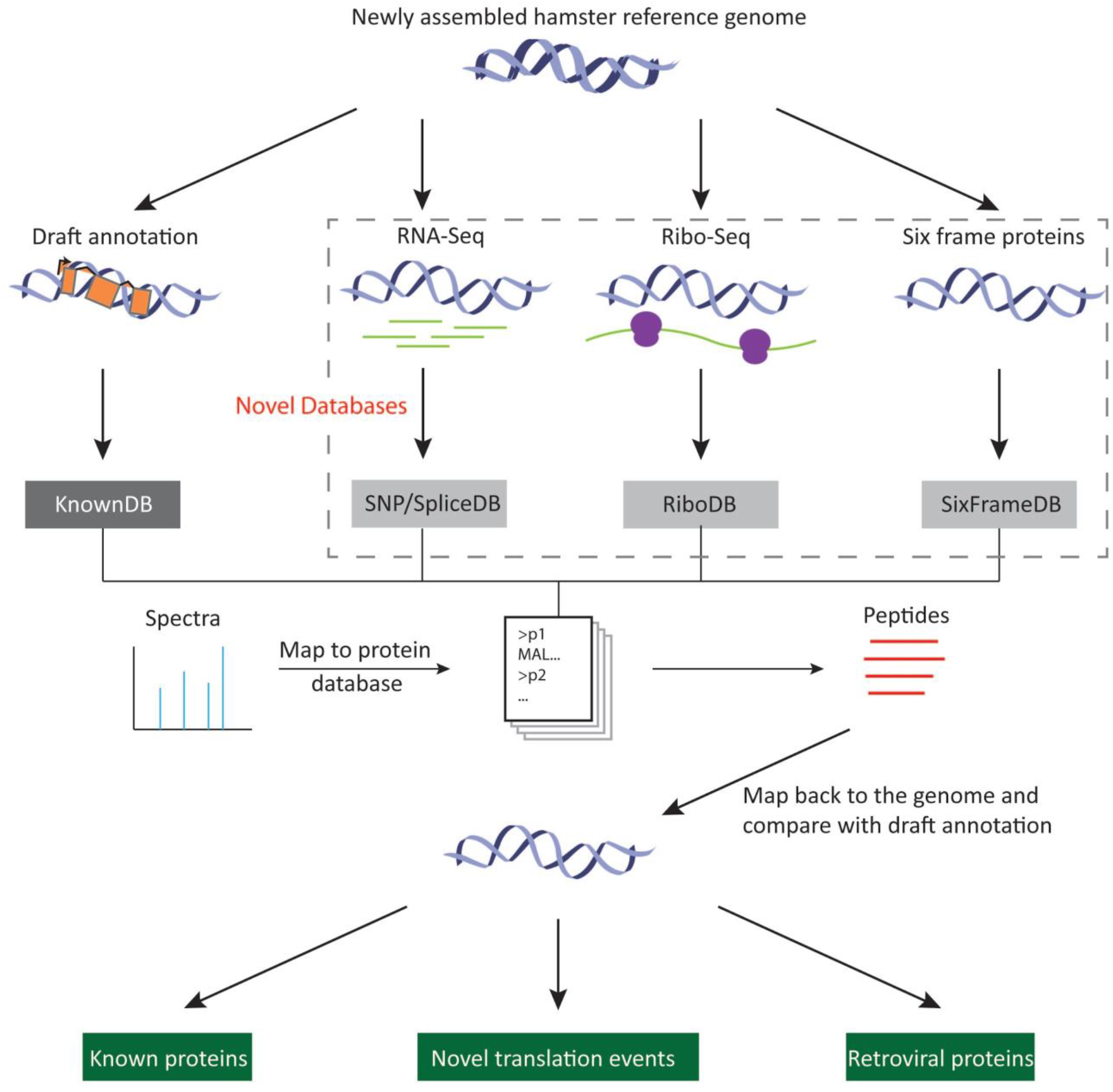
Overview of the proteogenomic pipeline. Multiple databases of putative protein sequences were generated based on the newly assembled hamster genome(Rupp et al., 2018) and additional data. The KnownDB contains protein sequences from our draft annotation generated here. The SNP/SPliceDB was derived from RNA-Seq samples, and contains candidate mutated or novel spliced proteins compared to draft annotation. The RiboDB was derived from predicted translated ORFs from Ribo-Seq and RNA-Seq. The SixFrameDB is derived from the reference genome(Rupp et al., 2018). After database construction, mass spectra were mapped against the protein databases using MSGF+ to identify the peptides. The peptides were then mapped back to the genome and compared with the draft annotation to verify translated known proteins, enumerate novel translation events and the identity of retroviral proteins.

## Results

### Draft annotation for the genome

The CHO-K1(Xu et al., 2011) and Chinese hamster genome sequences(Brinkrolf et al., 2013b; Lewis et al., 2013b) were originally assembled using short read (99bp) technology, and therefore resulted in fragmented contigs and scaffolds. Thus, efforts to annotate the genomes resulted in some errors in protein and gene models. The RefSeq pipeline has corrected some such errors; however, the complete reassembly of the Chinese hamster genome(Rupp et al., 2018) provides an opportunity to obtain a much improved annotation of coding regions in the genes and their corresponding protein sequences. Therefore, we generated a new draft annotation here **(Supplementary Figure S1)**, including predicted protein sequences.

68 RNA-Seq samples were prepared from multiple CHO cell lines and hamster tissues **(see Methods).** Transcripts were assembled for each sample separately using stringtie(Pertea et al., 2015) and merged using stringtie-merge(Pertea et al., 2015), yielding 26,530 genes with 68,082 transcripts. Then 38,654 hamster RefSeq transcripts were mapped to the newly assembled hamster reference genome using GMAP(Wu & Watanabe, 2005). Finally, the RefSeq alignments and RNA-Seq assembled transcripts were merged to 86,790 transcripts and grouped into 38,511 genes based on genomic locations using Program to Assemble Spliced Alignments (PASA).

We then applied TransDecoder(Haas et al., 2013) to the 86,790 transcripts and predicted 63,331 proteins with 47,829 unique protein sequences (the remaining are noncoding transcripts). To annotate the function of the proteins, we aligned the protein sequences to the hamster RefSeq and UniProt Swiss-Prot protein databases using BLASTP(Gish & States, 1993). **(Supplementary Table S1)**. We assigned UniProt gene names to the proteins in our draft annotation except for those that only map to hamster RefSeq proteins. Furthermore, we identified 4,640 non-coding transcripts by aligning the transcripts to the hamster RefSeq non-coding transcripts using BLASTN.

### Proteogenomics helps identify novel proteins in the draft annotation

To quantify novel proteins predicted in the draft annotation compared to the hamster Refseq proteins, we mapped 47,829 unique draft proteins to the RefSeq proteins using BLASTP. We classified the mappings into 5 main categories: (1) 15,787 perfectly mapped proteins; (2) 7,483 proteins mapping perfectly on only one end between the draft and RefSeq sequences; (3) 11,780 high quality mapped proteins (over 90% percentage of identity (pident) and over 80% percentage of length (plen) on both sides of homology protein mapping pair between draft and RefSeq); (4) 6,688 high quality mapped proteins (over 90% pident and over 80% plen on either side), but only mapping well on one end; (5) 5,820 low quality or non-mapping proteins. 289 proteins failed to map. We defined proteins that were not in category (1) as novel proteins. Among the one-sided perfect mapping proteins, 3,336 proteins are shorter in the draft, compared to RefSeq, while 4,147 proteins are longer than RefSeq. Interestingly, isoforms of some of the former proteins map perfectly to RefSeq, which indicates the draft annotation pipeline is sensitive to splice sites, which resulted in more isoforms being assembled than seen in Refseq.

Next, we sought peptide support for the novel proteins in the draft annotation. 12,870,725 mass spectra were acquired and prepared from multiple CHO cell lines and hamster tissues. We merged the draft and RefSeq proteins and extracted the unique protein sets as a reference protein database. Then we used MS-GF+(S. Kim & Pevzner, 2014) to search the peptide-spectrum matches (PSMs) with 1% FDR correction and identified 244,729 peptides. Here we only consider proteins with at least two peptides and at least one of them mapping uniquely. For each pair of homologous draft and RefSeq proteins, the draft protein was considered as novel if it has extra peptide support compared to the corresponding RefSeq protein. As a result, we identified 4,077 draft novel proteins, 3,890 of which have additional unique peptide support not seen in the RefSeq sequence (**Figure 2A**). The remaining 187 novel proteins have extra peptides mapping to multiple locations. **(Figure 2B)**. The high quality protein mappings category (>90% pident and >80% plen) has the most novel proteins. 5,608 draft proteins have the same peptide support as similar RefSeq proteins, which may require additional data to verify their novel features. The numbers of novel proteins in each mapping category are depicted in (**Figure 2)**.

**Figure 2:**
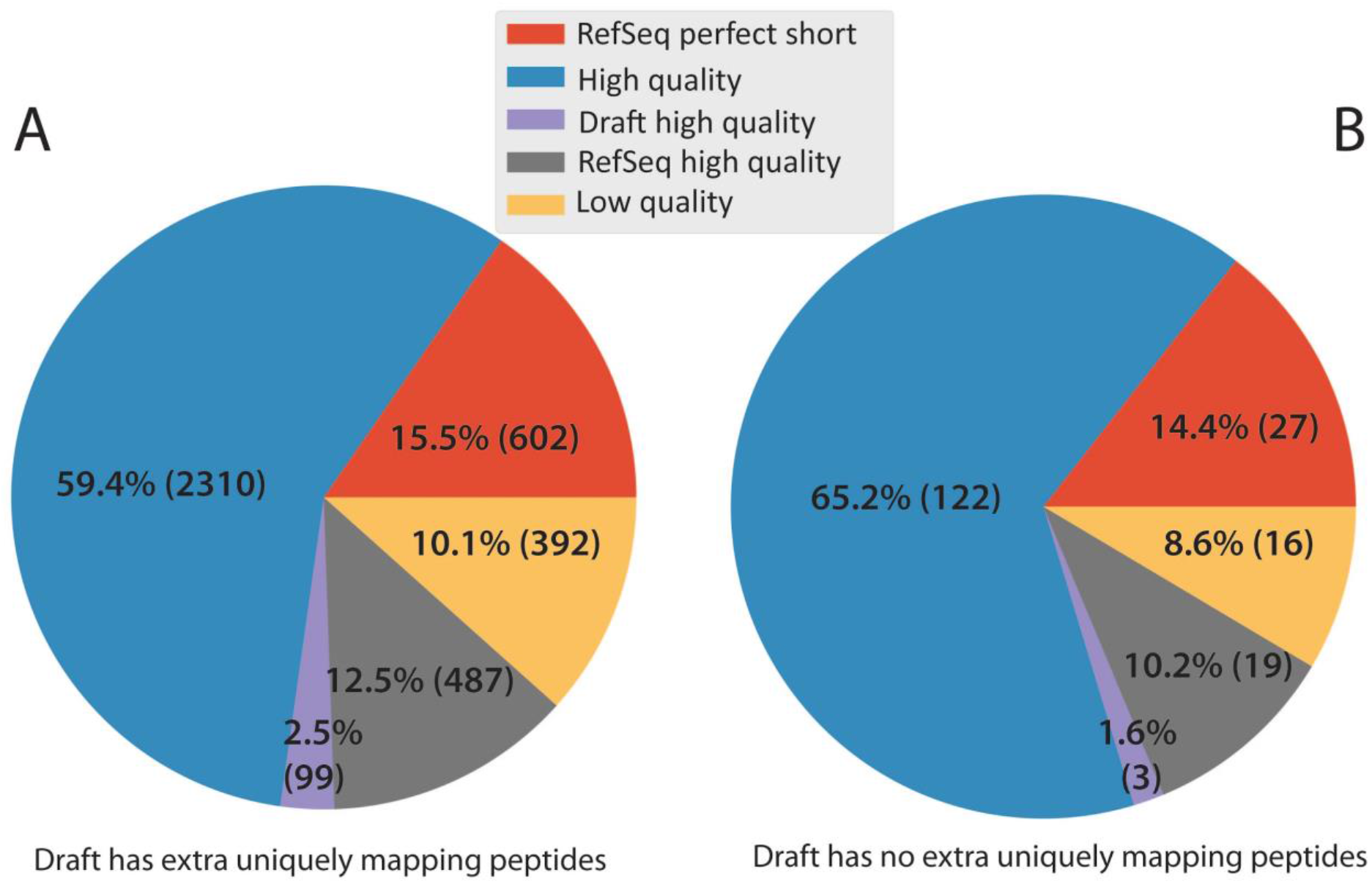
Number of novel draft proteins verified by draft-only peptides in different categories. The draft annotation predicted thousands of novel protein sequences. **(A)** Of these 3,890 had uniquely mapped peptides supporting the novel protein sequences. **(B)** Only 187 did not have extra peptide support from uniquely mapping peptides, and thousands provided peptide support. **RefSeq perfect short**: RefSeq proteins map perfectly but are shorter than draft proteins; **High quality**: high quality mapping proteins between draft and RefSeq; **Draft high quality**: draft proteins map to RefSeq with high quality, but the reverse doesn’t hold; **RefSeq high quality**: RefSeq proteins map to draft with high quality, but the reverse doesn’t hold; **Low quality**: low quality mapping between draft and RefSeq.

### Proteogenomics and Ribosome profiling identify additional translational events

In addition to verifying novel protein sequences, proteomics can also help identify other translational events, e.g., novel splice sites, gene fusions, etc. **(Figure 3A)**. Thus, to obtain a more comprehensive view of protein sequence verification and identification of other translation events, we created 4 putative protein databases: (1) KnownDB-predicted from draft annotation, (2) SnpDB-translated from RNA-Seq reads that have non-synonymous mutations and short INDELs, (3) SpliceDB-translated from spliced RNA-Seq reads, and (4) SixframeDB-peptides between stop codons in all 6 frames of the genome **(Figure 1)**. As previously recommended(Woo et al., 2015), we performed multi-stage 1% **(Supplementary Figure S2)** FDR correction for the databases sequentially. This pipeline **(Supplementary Figure S3)** identified 3,656,801 (28%) significant PSMs resulting in 239,973 unique peptides mapping to the KnownDB and 9,808 peptides mapping to the remaining databases **(Figure 3B)**. Among all the peptides identified from KnownDB, 208,904 (87%) map to unique genomic locations.

**Figure 3:**
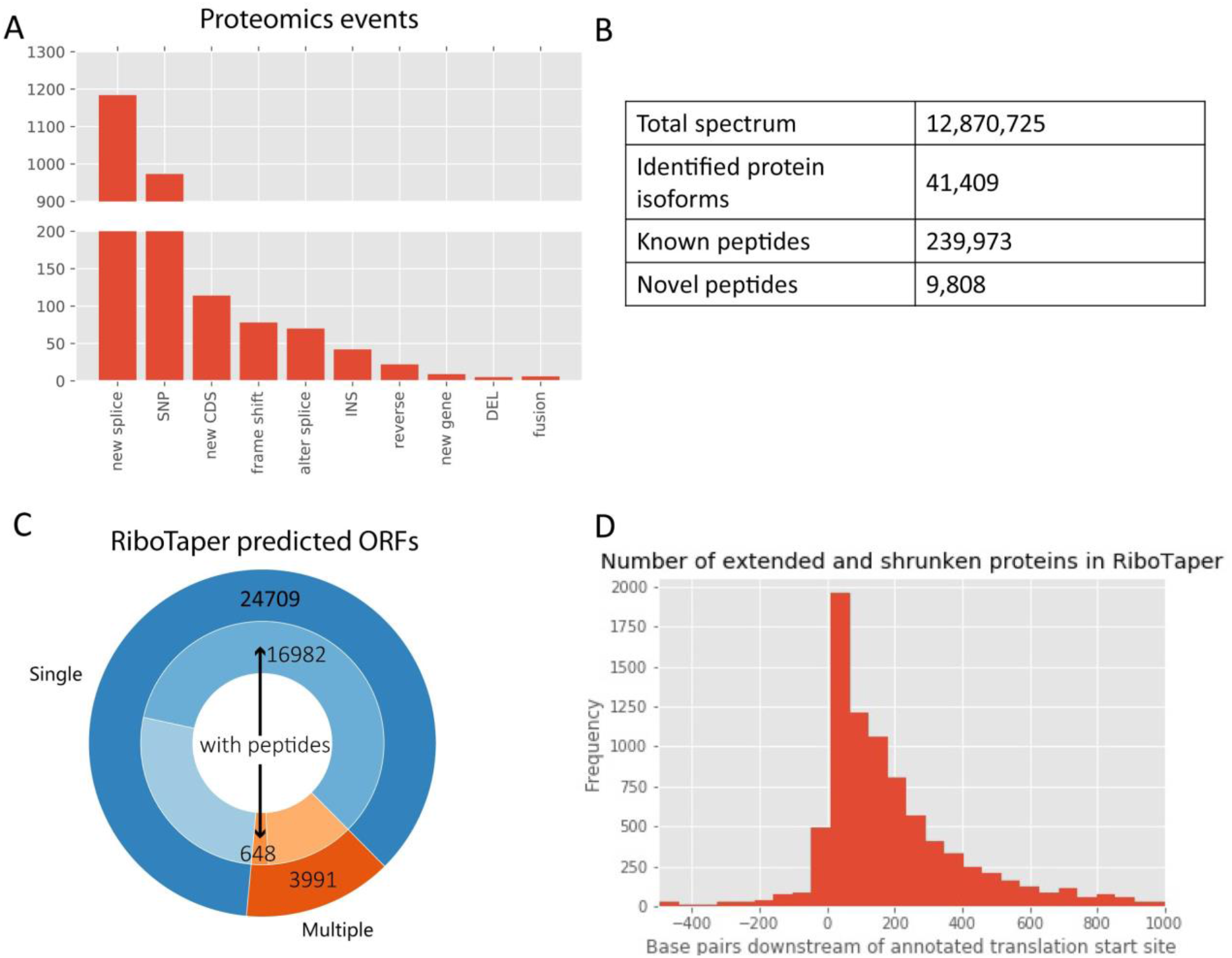
Proteogenomics and RiboTaper verified predicted protein sequences and identified novel translation events. **(A)** Numerous novel translational events were identified, including novel splice sites that are not in the draft annotation file (new splice), non-synonymous mutations (SNP), peptides that map to UTR regions or to transcripts with no CDS (new CDS), alternative splice sites (alter splice), peptide mapping to reverse strand of reference CDS (reverse), insertions (INS), peptide mapping to intergenic regions (new gene), deletion (DEL), and gene fusions connecting two genes (fusion). **(B)** Statistics for the number of spectra, peptides and protein isoforms identified in proteogenomics. **(C)** Number of ORFs identified using RiboTaper. **Outer circle:** Number of transcripts predicted with single ORF (blue) or multiple ORFs (orange). **Inner circle**: Number of transcripts with (darker blue and orange) or without (light blue and orange) peptide support. **(D)** Number of proteins that are shorter/longer than the draft annotation. Positive x axis means the RiboTaper proteins are shorter (i.e., start later) than the draft annotation.

We required each validated protein to have at least two peptides and at least one of them map uniquely (i.e., to only one locus). Using this strategy, we verified 30,814 proteins, which represent 64.4% of the sequences in the KnownDB.

After known protein sequence validation, we explored additional novel peptide events. To guarantee high confidence of the events, we filtered out those with fewer than 3 RNA-Seq supporting reads, and required each event to have at least one uniquely mapped peptide. Most proteins are longer than 100 amino acids, and tryptic peptides sequenced are usually about 7-32(Frank, 2009) amino acids long. Thus, if novel peptides are close and in the same translational frame, they are likely to support the same protein. Therefore, we clustered novel identified peptides that were in the same translational frame and fewer than 100 amino acids away from each other to represent the same event. In total, we discovered 2,503 new translational events, 86% of which are novel splice and nonsynonymous single nucleotide polymorphisms (SNPs) **(Figure 3A)**. Novel splice sites represent 47.3% of the total events, covering 857 genes. We also identified 70 alternative splice events, 22 reverse strand translation events, 9 novel ORFs (not predicted by the draft annotation) and 5 gene fusion events.

Ribo-Seq offers orthogonal evidence to further support protein sequences, and can help discover new proteins as well. Ribo-Seq and corresponding RNA-Seq data sets for an IgG-producing CHO cell line were acquired at both exponential and stationary phase(Kallehauge et al., 2017). We used RiboTaper(Calviello et al., 2016) to predict the translating ORFs under the guidance of the draft annotation. 28,700 transcripts were predicted to encode proteins, with 24,709 (86%) having a single ORF **(Figure 3C)**. Among these, 13,666 transcripts have the same protein sequences as the draft annotation. In addition, 1,318 “non-coding transcripts” in the draft annotation are predicted to encode proteins. The remaining Ribo-Seq predicted sequences were classified into two groups: (1) in the same frame (8775) and (2) in different frames (950) with the draft annotation **(Figure 3D, Supplementary Table S2)**.

Protein predicted by Ribo-Seq can expand the databases for proteomics so that more proteins can be verified. Here we used Ribo-Seq predicted proteins as a novel database and ran the proteogenomics pipeline together with KnownDB. After filtering all peptides identified in the previous proteogenomics pipeline, the Ribo-Seq data facilitated the identification of 2,581 new peptides. Here we require at least one unique peptide in an ORF to verify translation. Among 1,628 non-protein-coding transcripts in the draft annotation, 218 were verified to encode proteins by Ribo-Seq and proteomics. While Ribo-Seq enabled the successful identification of many new peptides, including those in transcripts previously thought to be non-coding, it was less successful at identifying the translation start sites. Supporting peptides were found for only 8 out of the 305 ORFs that were predicted to be longer than the draft annotation. In addition, Ribo-Seq helped identify translation events in 5’UTR and 3’UTR regions in 234 genes, consistent with previous reports in human(Ingolia et al., 2014). For more details, see **Supplementary Table S2.**

### Proteomic based validation of SNPs and INDELs in CHO cell lines and hamster tissues

The peptides obtained from proteomic studies enabled the validation of genetic variants in the various CHO cell lines and hamster tissues. To discover these mutations, we used our SnpDB **(Figure 1)** as the novel database in the proteogenomics pipeline, which includes peptides translated from all the RNA-Seq reads supporting SNPs and small INDELs. Proteomics identified fewer mutations than RNA-Seq **(Supplementary Table S3)**, mainly because of its lower depth of coverage compared to RNA-Seq. Furthermore, mutated proteins can be degraded, and therefore not detected. In total we identified 973 nonsynonymous SNPs, located in 734 genes. Most genes have one SNP while there are 6 genes with more than 5 SNPs: GAPDH, GOLGB1, AHNAK2, PKM, MYH9 and EEF1A1. Surprisingly, only one protein lost a stop codon (ribosome protein gene RPS23) while others change amino acids. 75% of the SNPs are homozygous, which indicates CHO cell lines may have developed and retained those mutations after long periods of evolution. Furthermore, the distributions of the 6 SNP types showed that transitions occurred more frequently than transversions, and the proteomic data captured a similar distribution of mutations as RNA-Seq data **(Figure 4A, Figure 4B).** We identified 42 insertions and 6 deletions, located in 41 and 6 genes respectively. There are more homogeneous frameshift INDELs than other types. The 8 genes harboring both SNPs and insertions include AHNAK2, CALR, HNRNPUL1, HSP90B1, PLEKHG5, PTMA, RIF1, and VAT1, while only SIK3 had both SNPs and deletions.

**Figure 4:**
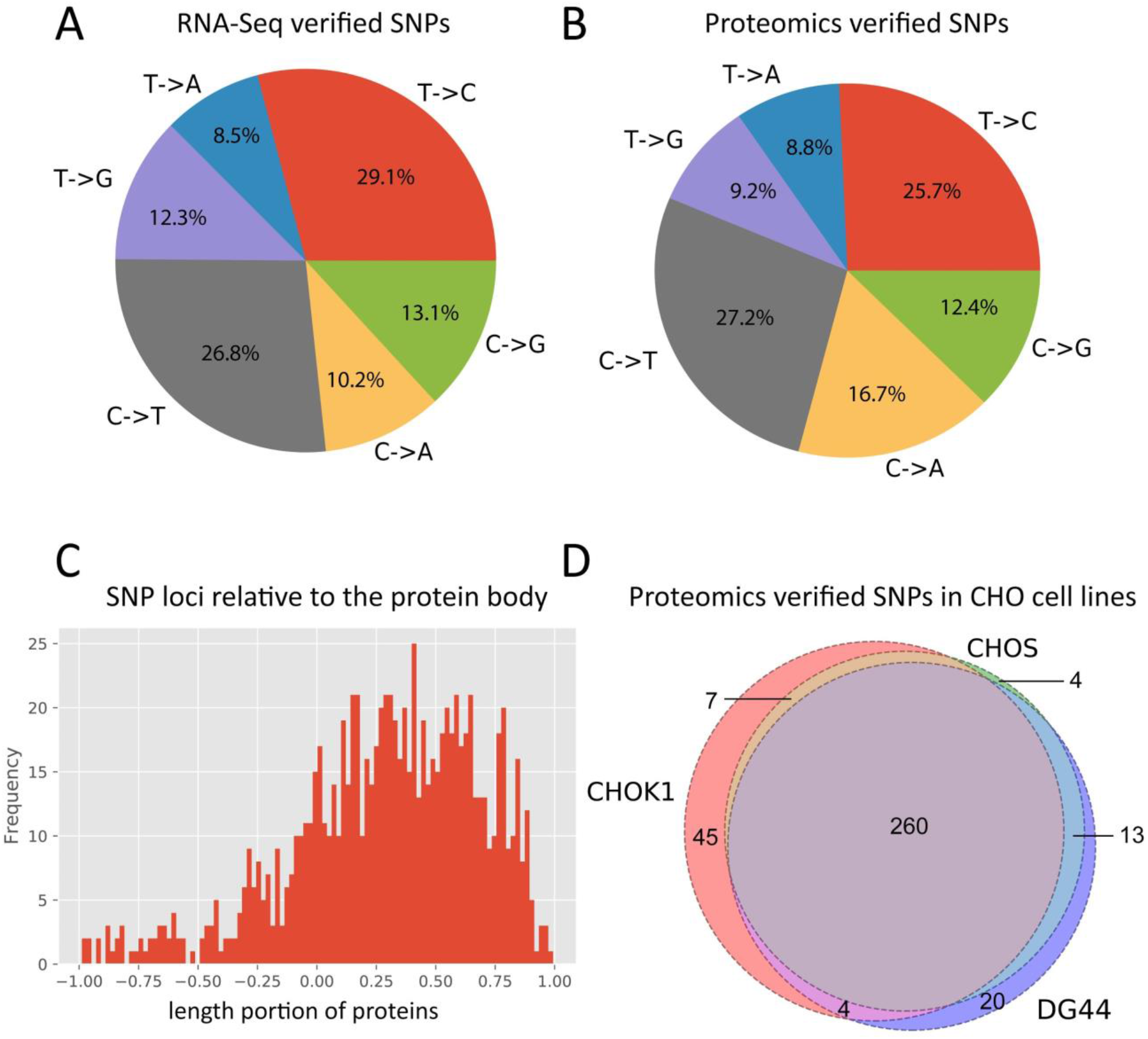
Hundreds of SNPs in hamster and different CHO cell lineages are validated. A comparison of the **(A)** distribution of SNP types identified from RNA-Seq and **(B)** SNP types verified by proteomics validates the overall distribution of SNPs. **(C)** Peptide-validated non-synonymous SNPs are located throughout the protein bodies. The length of each protein is scaled to 1. 0 represents the start codon. SNPs that locate below 0 or above 1 represent peptide-supported SNPs in 5’-UTR and 3’-UTR regions, respectively. **(D)** Venn diagram of 353 peptide-supported SNPs from CHO-K1, CHO-S and DG44 cell lines shows that most SNPs are shared across cell lines.

Next, we looked at the mutation distribution across the protein body. Since there are far fewer INDELs than SNPs, focused on the distribution of SNPs. **Figure 4C** shows that SNPs are distributed relatively evenly across the protein body. Interestingly, we also identified peptide-supported mutations in 5’ UTR regions, but the number is much smaller than for those in coding regions.

Many different CHO cell lines (e.g., CHO-K1, CHO-S and DG44), have been used to develop different recombinant protein-producing cells. Each cell line has a lengthy history of mutation and selection during cell line development(Lewis et al., 2013a), and therefore can have unique genomic variants(Lewis et al., 2013a; van Wijk et al., 2017). To check the variations between these CHO cell lines, we compared their peptide-supported SNPs in the coding regions. Most of the SNPs are shared among the cell lines, which means these variants have been conserved during the long period of cell line development, either due to early mutations obtained in CHO cells when derived in 1957 or genetic drift of the Chinese hamster colony since then **(Figure 4D)**.

### Proteomics elucidate translated retroviral elements in the genome

For decades, it has been known that CHO cells shed retroviral particles(Lieber, Benveniste, Livingston, & Todaro, 1973); while these were shown to be non infectious(Anderson, Low, Lie, Keller, & Dinowitz, 1991; Dinowitz et al., 1992), the safety concern has required companies to filter out all such viral particles and conduct extensive testing to verify non-infectivity. This adds a substantial cost to production. Viral particles have been isolated, but all mRNAs that have been sequenced from these particles were all non-coding, in that they contained many early stop codons. Thus, it remains unclear which loci encode the translated viral particles. Here, we analyzed the transcriptomic and proteomic data to identify the loci of expressed and translated retroviral particles, to enable further efforts to eliminate these particles and reduce drug purification costs.

To identify translated endogenous retroviruses in CHO cells, we first extracted peptides that support retroviral proteins from our draft annotation **(Figure 5A)**. Since retroviruses can have multiple similar copies across the whole genome, we consider peptides that can map to multiple locations. We found 457 retroviral genes covered by 723 transcripts, 184 of these genes (corresponding to 304 transcripts) have peptide support and 111 transcripts map well to RefSeq (>60% pident, >80% plen) and UniProt (>50% pident, >80% plen). Infectious retroviral DNA is flanked by two identical non-coding repeats called long terminal repeats (LTR)(Temin, 1982), which aid in retroviral mobility and integration into the host genome, along with regulation of retroviral gene expression. In the hamster genome, we identified 3,324 LTR pairs in the reference genome using LTRharvest(Ellinghaus, Kurtz, & Willhoeft, 2008) and found only 76 retroviral transcripts locate between those LTRs, 74 of which have peptide support. If one LTR is disrupted, the other side can still effectively induce transcription. Thus, genes not flanked by LTR pairs can still produce retroviral particles.

**Figure 5:**
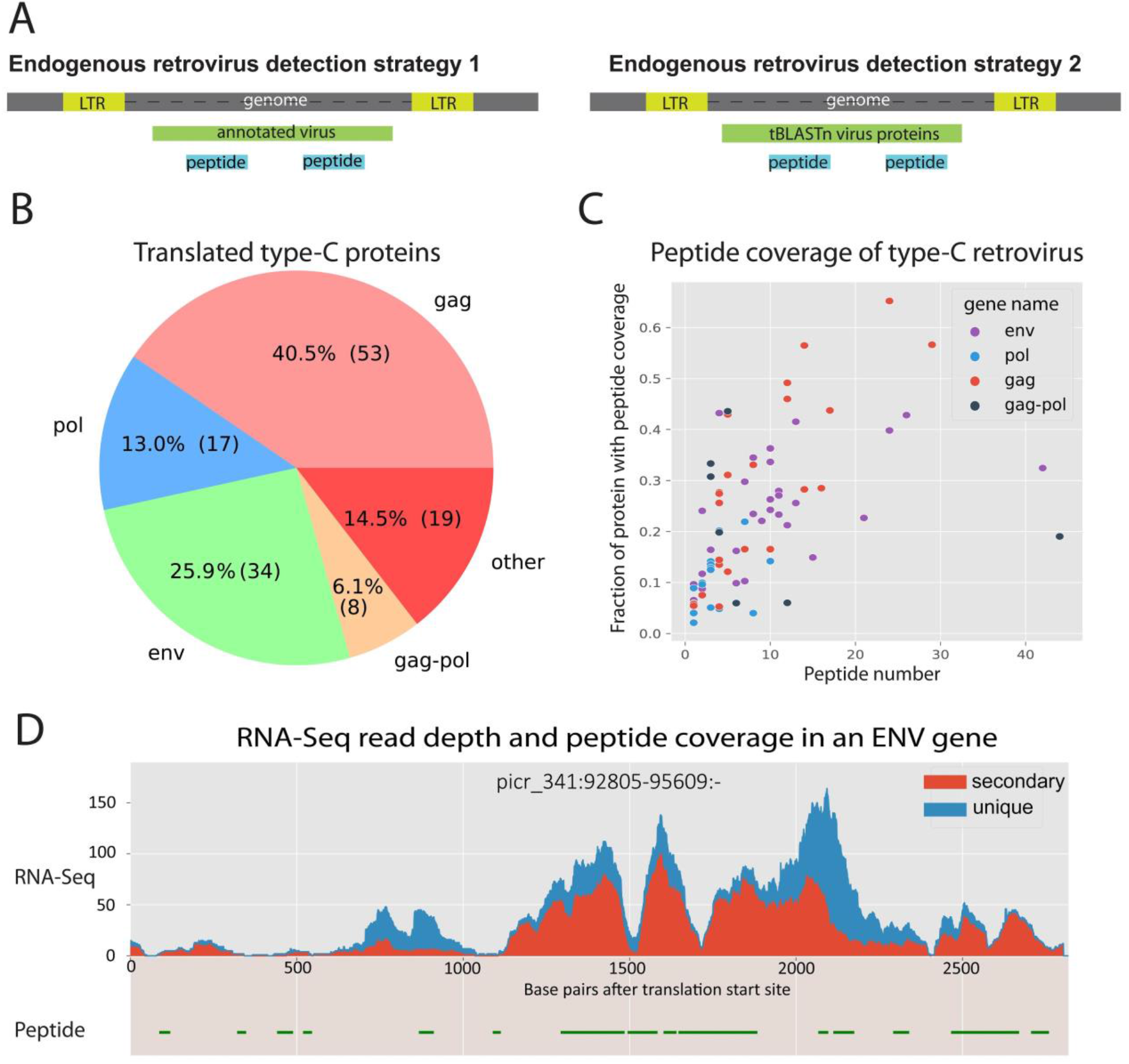
A proteogenomic identification of the source of translated endogenous retroviral particles shed from CHO cells. **(A)** Two strategies were taken to identify translated retroviral loci. In strategy 1, peptides were mapped to the annotated retroviral proteins. For strategy 2, the sequences from the NCBI retroviral protein database were aligned to the genome using BLASTP. Then we evaluated the overlap of these aligned peptides with the novel peptides identified from the novel databases in our proteogenomics pipeline. **(B)** The strategies recovered 131 type-C peptide-supported retroviral proteins in CHO cell lines (the “other” category represents non-typical retroviral proteins, such as the p12 protein). **(C)** Peptide-supported type-C virus proteins were analyzed to assess the portion of protein sequence covered by peptides against peptide number. **(D)** Coverage of an envelope protein in reverse strand. **uni**: reads map uniquely to the locus, **sec**: reads are secondary reads and map to multiple loci.

We next aimed to identify unannotated translated retroviral sites in the genome **(Figure 5A)**. For this, we aligned all retroviral proteins from NCBI to the reference genome using tblastn(Altschul, Gish, Miller, Myers, & Lipman, 1990). Then we overlapped the mapping sites with the 5,104 novel peptides identified against novel protein databases from our proteogenomics pipeline and obtained 41 novel retroviral sites, 1 of which localized between an LTR pair and was covered by 5 peptides. The site showed homology to the gag proteins from Gibbon ape leukemia virus and Spleen focus-forming virus. Finally, since CHO cells were originally derived in 1957, we further checked if new infections may have emerged in CHO (**Supplementary Figure S4).** For this analysis, we aligned all known retroviral proteins and novel peptides from proteogenomics pipeline, using tblastn, to in-house Illumina-corrected single molecule real time (SMRT) sequence data from CHO-S and the Chinese hamster(Rupp et al., 2018). After filtering out the putative viral sites identified in hamster, we found no evidence that wild type CHO-S cell lines have acquired any new retroviral sites.

Mammalian retroviruses have been classified into different types based on the genomic compositions. We mapped retroviral protein sequences identified from previous steps to UniProt and used protein full names to discover 3 main retroviral types in hamster: type-A, type-B, and type-C(Weiss, 1996). We found type-C retroviruses to be the most highly transcribed and translated **(Supplementary Figure S5)**. In addition, type-C viral particles have been identified in CHO cell lines before and regulatory agencies now require the verification that products are non-infectious type-C particles(Dinowitz et al., 1992; Lie et al., 1994). The vast majority of type-C proteins had little or no peptide support, suggesting these are silenced or noncoding. Of the 131 type-C retroviral proteins with peptide support that we identified, most were gag and envelope proteins **(Figure 5B)**. Only 7 proteins had more than 20 peptides, and most proteins had low coverage of supporting peptides **(Figure 5C). Figure 5D** shows an example of highly translated envelope protein with 42 peptides, covering 32% of the coding sequence. Although it has many secondary reads, it also has many uniquely mapping reads, which indicates this locus is truly expressed. The proteins with high coverage and more supported peptides should be prioritized in efforts to eliminate viral particle production. The genomic locations, RNA-Seq and peptide coverage of all endogenous retroviral genes are provided in **Supplementary Table S4**.

## Discussion and Conclusion

Here we presented the first proteogenomic reannotation of the Chinese hamster genome, in which we utilized RNA-Seq, Ribo-Seq, and proteomics to improve the annotation. To identify as many peptide-supported proteins as possible, we mapped spectra to a known protein database from a draft annotation and several novel protein databases derived from different data types. We identified 4,077 novel proteins in the draft annotation compared to the hamster RefSeq protein database and 2,503 novel translational events and mutations in hamster and CHO cell lines. Furthermore, we identified the potential sources of retroviral particles shed from CHO cells, including 131 type-C retrovirus genes, 7 of which are supported by more than twenty peptides.

Usually an annotation is required before running proteogenomics pipeline. The typical first step of genome annotation is masking the repeats to avoid getting millions of seeds during BLAST(Yandell & Ence, 2012). However masking the genome can hide important annotation information, including common domains and retroviral elements. To avoid this loss of information, we aligned assembled transcripts to the unmasked genome using gmap. Doing so resulted in only 0.7% transcript mappings to be ambiguous (i.e., with >2 mappings). This enabled the annotation and analysis of endogenous retroviral genes, which usually have multiple similar copies. Masking repeats often removes such information(Slotkin & Keith Slotkin, 2018). Thus, future annotation efforts would benefit from the acquisition and alignment of high quality transcripts or protein sequences to the genome of interest.

Different pipelines have been designed for genome annotation in higher eukaryotes(Yandell & Ence, 2012), and most include *ab initio* prediction, mapping of homologous protein sequences, and transcript assembly and alignment. As the throughput and resolution of mass spectrometry techniques is increasing, more studies are integrating these data type to refine annotations(Nesvizhskii, 2014). Here we discovered thousands of genes and novel translational events using proteomics data. Despite their value, proteomics data can be sparse since the chemical properties of some peptides are less compatible with the experimental setup (e.g., due to hydrophobicity) or size of the peptides post digestion(Chandramouli & Qian, 2009). Furthermore, many peptides have diverse post-translational modifications, thus making it difficult to align peptides to predicted peptide sequences from genomes. Thus, Ribo-Seq provides a complementary method to further discover new genes or correct annotations(Ingolia, Brar, Rouskin, McGeachy, & Weissman, 2013). The Ribo-Seq datasets we used here are from the DG44 CHO cell line, and CHO cells may only express half to two thirds of their genes(Rupp et al., 2014; Singh, Kildegaard, & Andersen, 2018). Thus, further annotation efforts would benefit from the acquisition of Ribo-Seq from many other hamster tissues and developmental stages. In summary, as more tools and proteomic and Ribo-Seq data accumulates, these data types will become increasingly integrated into standard pipelines for genome annotation.

Finally, the reannotation here provides an invaluable resource for the development of improved CHO cell lines. While CHO cells provide several advantages as an expression host for recombinant protein production, HCPs are continuously secreted(Hogwood, Bracewell, & Smales, 2014; Kumar et al., 2015), thus impacting recombinant protein quality and safety. Thus, expensive chromatographic columns, filtration systems, and infection and HCP assays are required during downstream processing. This adds considerable cost to biopharmaceuticals. One particular regulatory concern has been the endogenous retroviruses that are shed by CHO cells(Lie et al., 1994). For these, assays were developed to quantify retroviral particles and ensure infectious retroviral particles are not found in the drug product after extensive filtration(de Wit, Fautz, & Xu, 2000). Here, we identified, among hundreds of retroviral genes in the hamster genome, which ones are expressed and translated in several CHO cell lines. This information will enable future efforts to remove these from CHO cells, thereby reducing burdens to downstream processing. To further facilitate such efforts, many endogenous retroviral genes show high levels of homology. In our work, this was manifested in the identification of many retroviral RNA-Seq reads and tryptic peptides that map to multiple genomic loci. As many of these also share DNA sequence, multiple viruses can be knocked out simultaneously by targeting the conserved regions, as accomplished in the pig(Yang et al., 2015). Doing this in CHO cells can reduce the costs by simplifying expensive purification steps where product can also be lost, and also simplify viral testing steps for the final product.

In conclusion, our work provides a refined and more extensive annotation of the Chinese hamster genome, which will enables more accurate CHO cell line engineering(Fischer, Handrick, & Otte, 2015; Kuo et al., 2017; Lee, Grav, Lewis, & Faustrup Kildegaard, 2015; Richelle & Lewis, 2017)The improved annotation will also facilitate improved processing of omics data(Chen, Le, & Goudar, 2017) as the gene models improve. Finally, a more complete list of all genes will enable efforts to map out molecular pathways in CHO cells to enable systems approaches to cell line development(Kuo et al., 2017).

## Methods

### Proteomic sample preparation

Proteomic data were acquired at two different locations using different protocols, thus increasing the diversity in spectra used for annotation. These are referred to as batch 1 and batch 2, as follow.

### Tissue Sample Collection for batch 1

Chinese hamsters were generously provided by Dr. George Yerganian (Cytogen Research, Roxbury, MA). Euthanization was performed by CO_2_ and verified by puncture. Harvested liver and ovary tissues were flash frozen on dry ice and stored at −80°C until analysis.

### Cell Culture Sample Collection for batch 1

Suspension CHO cell lines (including CHO-S and CHO DG44) were grown in shake-flask batch culture. CHO-S cells were cultured in CD-CHO medium supplemented with 8mM glutamine (Thermo Fisher Scientific, Waltham, MA), and CHO DG44 cells were culture in DG44 medium supplemented with 2mM glutamine (Thermo Fisher Scientific, Waltham, MA). Samples were collected on day 2 for exponential phase and day 4/5 for stationary phase. Cells were incubated at 37°C, 8% CO_2_, and 120RPM. For sample collection, approximately 3 million cells were spun down, washed with PBS on ice, frozen rapidly on dry ice, and stored at −80°C until analysis.

### Cell lysate and Tissue Sample preparation for batch 1

Cell culture lysates and tissue samples were thawed on ice and suspended in 2% sodium dodecyl sulfate (SDS) supplemented with 0.1mM phenylmethane sulfonyl fluoride (PMSF) and 1mM ethylenediaminetetraacetic acid (EDTA), pH 7-8. Samples were lysed by sonicating for 60 seconds at 20% amplitude followed by 90 seconds pause (for three cycles). Protein concentration was measured with bicinchoninic acid (BCA) protein assay after briefly spinning to remove cell debris. Three hundred micrograms of each sample was reduced in 10mM tris(2-carboxyethyl)phosphine (TCEP), pH 7-8, at 60°C for 1hr on a shaking platform. After bringing each sample to room temperature, iodacetamide was added to alkylate the sample to 17mM final concentration for 30 minutes. Next, samples were cleaned using 10kDa filters to reduce the SDS concentration as suggested by the filter aided sample preparation (FASP) protocol(Wiśniewski, Zougman, Nagaraj, & Mann, 2009). The samples were finally digested using trypsin/LysC enzyme mix at an enzyme to substrate ratio of 1:10 (Promega V507A, Madison, WI), overnight at 37°C on a shaking platform.

### Identification of Proteins by Mass Spectrometry for batch 1

Digested peptides (100µg from each protein digest) were fractionated on a basic reversed phase column (XBridge C18 Guard Column, Waters, Milford, MA). Fractions were concatenated into 48 prior to second dimension LC and MS analysis. The use of fractionation with equal peptides in each was designed to mimic biological replicates for each sample. Tandem MS/MS analysis of the peptides was carried out on the LTQ Orbitrap Velos (Thermo Fisher Scientific, Waltham, MA) MS interfaced to the Eksigent nanoflow liquid chromatography system (Eksigent, Dublin, CA) with the Agilent 1100 auto sampler (Agilent Technologies, Santa Clara, CA). Peptides were enriched on a 2cm trap column (YMC, Kyoto, Japan), fractionated on Magic C18 AQ, 5µm, 100Å, 75µm x 15cm column (Bruker, Billerica, MA), and electrosprayed through a 15µm emitter (SIS, Ringoes, NY). Reversed phase solvent gradient consisted of solvent A (0.1% formic acid) with increasing levels of solvent B (0.1% formic acid, 90% acetonitrile) over a period of 90 minutes. LTQ Orbitrap Velos parameters included 2.0kV spray voltage, full MS survey scan range of 350-1800m/z, data dependent HCD MS/MS analysis of top 10 precursors with minimum signal of 2000, isolation width of 1.9, 30s dynamic exclusion limit and normalized collision energy of 35. Precursor and fragment ions were analyzed at 60000 and 7500 resolutions, respectively.

### In-solution digestion of whole cell lysate for proteomics batch 2

A second batch of samples were prepared and analyzed using a different approach. For these, 1mg of protein sample was transferred to a centrifuge tube, and all samples were equalized to the same volume using the same lysis buffer. A fresh stock of 0.5M reducing agent dithiothreitol (DTT) was prepared, and an appropriate volume of DTT was added to achieve a final concentration of 5mM. Samples were incubated for 25 minutes at 56°C. Before alkylation samples were cooled to room temperature and an appropriate volume of freshly prepared 0.5M iodoacetamide was added to a final concentration of 14mM and incubated for 30 minutes at room temperature. Untreated iodoacetamide was quenched by a second addition of 0.5M DTT to make total concentration of DTT equal to 10mM and incubated for 15 min at room temperature. The protein mixture was diluted 1:5 in 25 mM Tris-HCl, pH 8.2, to reduce the concentration of urea to 1.6 M. A double digestion by trypsin was performed by adding trypsin to 1/50 enzyme: substrate ratio and incubated at 37°C. After 4 hours of primary incubation the trypsin was topped up (enzyme: substrate ratio 1/100), and the protein mixture was left to digest overnight at 37°C. After the overnight digestion, unused trypsin was quenched by adding TFA to a final concentration of 0.4%.

### Peptide sample clean-up for proteomics batch 2

Digested peptides were desalted and cleaned up using Sep-Pak c18 Vac cartridge, 200mg sorbent per cartridge, 55-500 µm Particle size (WAT054945) using negative pressure. The C18 cartridge was washed and conditioned by using 9ml of ACN followed by 3ml of 50%ACN and 0.5% acetic acid. C18 resin was then equilibrated with 9ml of 0.1% TFA and samples were loaded in 0.4%TFA. Loaded samples were desalted with 9ml 0.1%TFA. TFA was removed with 1ml 0.5% acetic acid. Desalted peptides were eluted with 3ml of 50%ACN 0.5% acetic acid. The eluted fraction was applied twice and collected in a 15ml conical tube. The eluate was snap frozen in liquid nitrogen, and lyophilized overnight or until the white (sometimes yellow) fluffy powder was observed. Dried peptides were stored at −20°C or otherwise dissolved in the appropriate buffer for phosphopeptide enrichment.

### Draft genome annotation generation

68 RNA-Seq samples from multiple CHO cell lines and hamster tissues were trimmed and aligned to the newly assembled hamster reference genome using Trimmomatic 0.36(Bolger, Lohse, & Usadel, 2014) and STAR 2.5.2(Dobin et al., 2013), respectively. The aligned reads were assembled into transcripts using stringtie(Pertea et al., 2015) for each sample and then the transcripts were consolidated into a union transcript set using stringtie-merge(Pertea et al., 2015). To improve the transcript coverage, we also mapped the hamster RefSeq transcript sequences to the newly assembled hamster genome using gmap(Wu & Watanabe, 2005) which were then integrated with transcripts generated from stringtie-merge using the PASA pipeline(Haas et al., 2003). Potential proteins encoded in the transcripts were predicted using transdecoder(Haas et al., 2013). Finally the functions of predicted proteins were determined by mapping them to the hamster RefSeq proteins and UniProt Swiss-Prot proteins(The UniProt Consortium, 2018) using BLASTP(Gish & States, 1993). Proteins whose mapping lengths were greater than 80% were considered and percentage identity of 60% when mapping to hamster RefSeq and of 50% when mapping to UniProt were used as threshold to further classify proteins into 4 categories with 1 to 4 indicating decreasing confidence scores as follows: 1. Pident and plen are larger than the threshold and have the same gene name between hamster RefSeq and UniProt. 2. Pident and plen are larger than the threshold and have different names between hamster RefSeq and UniProt. 3. Pident and plen are less than the threshold and have the same gene name between hamster RefSeq and UniProt. 4. Pident and plen are less than the threshold and have different gene names between hamster RefSeq and UniProt. LncRNAs were predicted by aligned transcripts to hamster lncRNAs using BLASTN with pident larger than 60% and plen larger than 80%.

### Proteogenomics database construction

We prepared 4 protein databases for mass spectrum matching: known protein database (KnownDB), SNP database (SnpDB), splice database (SpliceDB) and six-frame translation of the genome database (SixframeDB). The KnownDB includes protein sequences extracted from draft annotation, the rest serve as novel protein databases. SnpDB was constructed by translating RNA-Seq reads that have mutations(Woo, Cha, Na, et al., 2014). Mutations were called using GATK3.7(Van der Auwera et al., 2013) and annotated using Annovar(Wang, Li, & Hakonarson, 2010). The SpliceDB was constructed by translating all RNA-Seq reads that span splice junctions(Woo, Cha, Merrihew, et al., 2014). The SixframeDB was derived from peptides fragments between stop codons in all frames of the reference genome assembly.

Peptide identification The original MS/MS spectra were converted from RAW format to Mascot Generic Format (MGF) using msconvert(Kessner, Chambers, Burke, Agus, & Mallick, 2008) and searched against each database independently using MSGFplus(S. Kim & Pevzner, 2014). Since different databases have different false discovery rates, it is recommended to perform multistage FDR correction with 1% cut off for the databases(Woo et al., 2015), which means the spectra failed to pass FDR correction were fed to the next database to correct again. We corrected FDR for databases in the following order: KnownDB, SnpDB, SpliceDb, SixframeDB.

### New translational event prediction

Significant peptide-spectrum matches (PSM) against the novel databases were used to discover new translation events using Enosi pipeline(Woo, Cha, Merrihew, et al., 2014). Briefly, identified novel peptides were mapped to the novel databases to get loci relative to their mapped proteins. Protein headers in the novel databases have loci relative to the reference genome assembly. A custom python script was used to deduce peptide loci relative to the reference genome assembly from those two loci. Then the loci were compared with the draft annotation to decide event type. Peptides in the same translation frame that are less than 300 nucleotides (100 amino acids) away are grouped to represent the same event. For SNPs and short INDELs, we filtered the false positives by variant calling using Illumina reads from sequencing hamster genomic DNA against the reference genome assembly.

### Protein prediction using Ribo-Seq data

We used previously published Ribo-Seq data of the CHO CS CS13-1.0 cell line(Kallehauge et al., 2017). All Ribo-Seq and RNA-seq data of each biological sample were trimmed and aligned to the reference genome assembly using Trimmomatic 0.36(Bolger et al., 2014) and HISAT 2.2.1(D. Kim, Langmead, & Salzberg, 2015) respectively. The aligned bam files were sorted and merged into one Ribo-Seq and one RNA-Seq bam files using Samtools 1.6(Li et al., 2009). The potential translated regions were predicted using RiboTaper(Calviello et al., 2016), which takes advantage of coverage in coding regions and triplet periodicity of ribosomal footprints. A custom python script was used to compare the translation prediction and draft annotation.

We treated the predicted proteins from Ribo-Seq as a novel protein database and combined it with the KnownDB. Then we mapped all the mass spectra to these two databases using the proteogenomics pipeline. In this case, the identified novel peptides would be the unique peptides from Ribo-Seq database.

### Retrovirus in the draft annotation

All viral proteins in draft annotation were identified by mapping to UniProt and hamster Refseq proteins using BLASTP and then the retroviral proteins were identify based on the full gene names. Peptides supporting annotated retroviral proteins were identified by mapping the known peptides to the KnownDB. LTRs were predicted using LTRharvest(Ellinghaus et al., 2008). Retroviral proteins that locate between LTR were identified by overlapping LTR regions with retroviral annotations using Bedtools2.27(Quinlan & Hall, 2010).

### New retrovirus discovery

Since virus proteins lack introns, we filtered novel peptides by removing those with splice sites. Retroviral proteins and filtered novel peptides were aligned to the reference genome assembly using tblastn(Altschul et al., 1990) and mappings with more than 60 pident and 55 plen were considered for downstream analysis. Virus mapping and peptide mapping were overlapped using Bedtools(Quinlan & Hall, 2010) to get the virus sites with peptide support, which were then further overlapped with LTR regions to get virus sites with both LTR and peptides support.

We decided the thresholds for virus tblastn mapping as follows **(Supplementary Figure S6)**. First, we started with low thresholds (30% for each). Then we assessed the overlap between filtered mapping and the draft annotation to identify those associated with known virus genes. If the mappings overlap with many non-viral genes, thresholds were incrementally increased and overlapped with draft annotation again. This was repeated until the mappings overlap with few non-viral genes and the number did not decrease anymore.

### Discovery of unique retroviruses in the CHO-S cell line

The in-house CHO-S SMRT sequence data was used to check if CHO-S has unique retroviral elements, compared to hamster. Firstly, we subsampled hamster SMRT reads to the same depth as CHO-S SMRT data, and both datasets were corrected using Illumina paired-end reads(Lewis et al., 2013a) through LoRDEC(Salmela & Rivals, 2014). Secondly, retroviral proteins and filtered novel peptides from our proteogenomics pipeline were mapped to corrected CHO-S and hamster SMRT reads using the same threshold as the previous tblastn mapping. Thirdly, virus and peptide mappings were overlapped to get virus sites with peptide support for CHO-S and hamster separately. Fourthly, we filtered CHO-S virus sites with hamster SMRT virus sites and mapped the unique CHO-S sites to the reference genome assembly. The mapped sites are the new retroviral sites and the unmapped sites are unique retroviral elements in CHO-S.

#### Type-C retrovirus detection in CHO cell lines

The functions of all the identified retroviral proteins were determined by mapping the protein sequences to UniProt using BLASTP. The full function descriptions of the proteins have the organism resource. Therefore, the types of all the retroviruses were determined by manually matching their full virus names to the types defined previously (see appendix “Retroviral Taxonomy, Protein Structures, Sequences, and Genetic Maps” of this book(Coffin, Hughes, & Varmus, 1997)). Peptide coverage of a protein equals to the number of amino acids covered by peptides divided by the protein length. The RNA-Seq coverage along the protein body was calculated using pysam(“Website,” n.d.).

## Accession

RNA-Seq raw data: PRJNA504034; Proteomics raw data in MassIVE: doi:10.25345/C5M597, Identified peptides in Synapse: doi:10.7303/syn17037373.

## Acknowledgements

This work was supported by generous funding from the Novo Nordisk Foundation provided to the Center for Biosustainability at the Technical University of Denmark (grant no. NNF16CC0021858), and SL was supported with funding from the Frontiers of Innovation Scholars Program at UCSD. VB was supported in part by grants from the NIH 1R01GM114362, and P-41-RR24851.

